# bacNeo: a computational toolkit for bacteria-derived neoantigen identification

**DOI:** 10.1101/2025.07.23.666275

**Authors:** Yunzhe Wang, Li Yuan, Katie W. Gu, Xiangdong Cheng, Bo Wen, Zhao Zhang

**Author notes:** Correspondence Author: Zhao Zhang.

## Abstract

Intratumour microbiome can restore anti-tumour immunity with bacterial peptides potentially serving as neoantigens, while robust *in-silico* tools for bacterial neoantigen identification remain lacking. We developed <u>bac</u>terial <u>neo</u>antigen (*bacNeo*), a multi-omics based computational toolkit that contains three modules to classify bacterial taxonomy, identify host HLA alleles, and predict bacterial neoantigens. *bacNeo* could enable personalised neoantigen discovery to advance therapeutic strategies across diverse cancer types.

## Main

As an emerging hallmark of cancer, the polymorphic microbiome can impact cancer phenotypes by mimicking tumorigenic signals, contributing to metastatic processes, and shaping the tumour immune microenvironment (TIME)^1^. As exogenous entities, intratumour microbes possess inherent immunogenicity that can reactivate immune responses^2^. For example, Kalaora et al. identified bacterial peptides presented by melanoma cells that elicited immune reactivity^3^. The processes started by neoantigen presentation via human leukocyte antigens (HLAs), which can subsequently be recognized by infiltrating T cells within the TIME, thereby reactivating antitumour immune responses^4^. However, the limited development of bioinformatics tools specifically designed to identify bacteria-derived neoantigens has hindered comprehensive investigations into their mechanistic roles and clinical applications in human cancers^5^.

Hence, we developed <u>bac</u>terial <u>neo</u>antigen (*bacNeo*), an intuitive and user-friendly bioinformatics toolkit that utilizes multi-omics data to identify bacteria-derived neoantigens with minimal installation requirements. *bacNeo* comprises three core functional modules: <u>bac</u>terial <u>c</u>omponents (*BACC*) for analysing bacterial taxonomies and phylogenesis; <u>bac</u>teria-bond <u>H</u>LAs (*BACH*) for determining sample-specific HLA alleles; and <u>bac</u>terial <u>p</u>eptides (*BACP*) for predicting HLA presented neoantigens from bacteria-derived peptides (Fig. 1).

**Figure 1.**
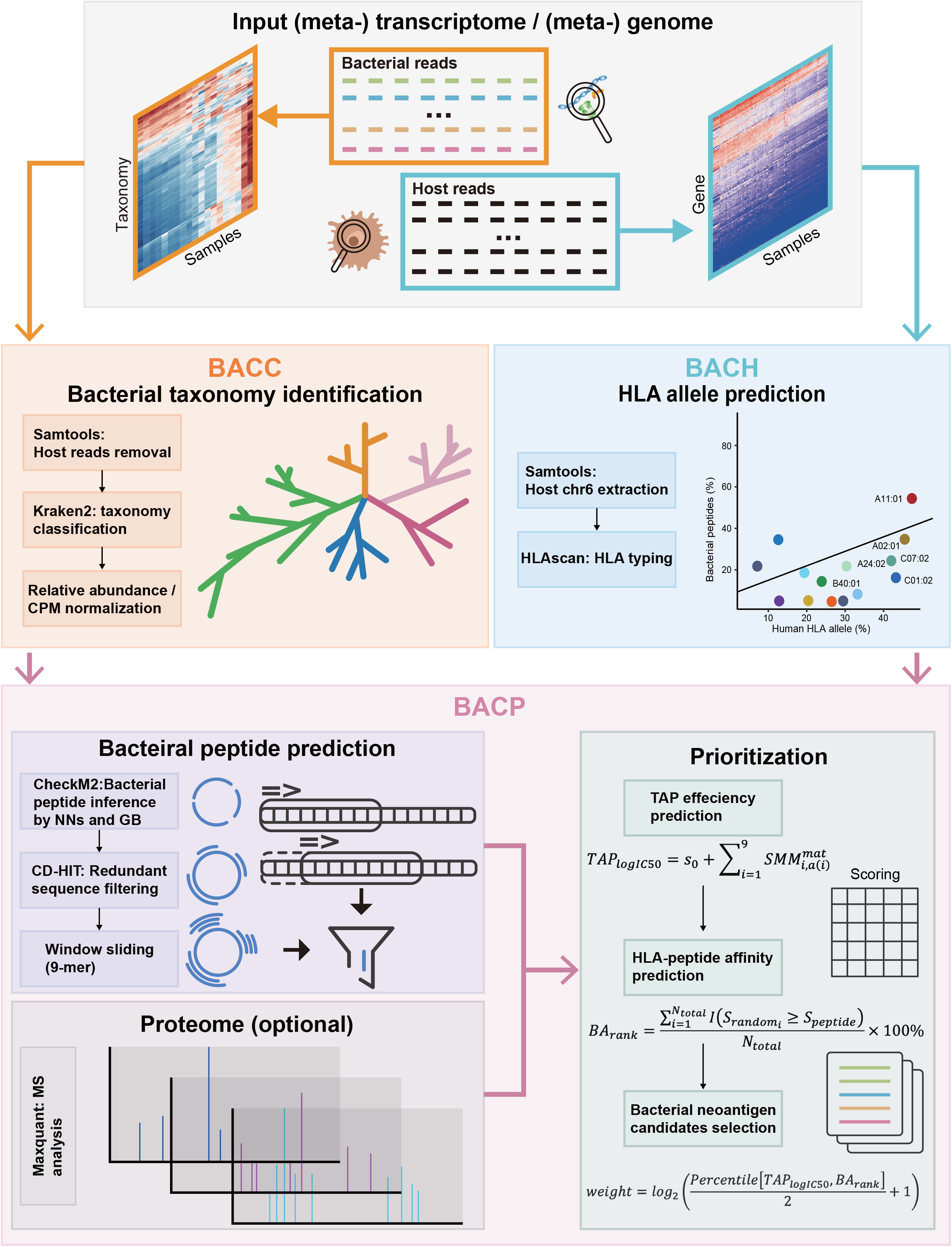
Overview of *bacNeo*, a bioinformatics tool predicting bacterial neoantigen candidates. *bacNeo* workflow illustrating data input and module interactions. Required inputs include sequencing data, with optional mass spectrometry data for enhanced validation. Arrows colour are paired with module-specific border colours.

The *BACC* module integrates established tools including BWA, HISAT2, and Kraken2^6^ to identify bacterial taxonomies from raw genomic (WGS/WES, recommended) or transcriptomic (RNA-seq) data. Initially, *BACC* utilizes BWA or HISAT2 to align sequencing reads to the host genome or transcriptome. Reads that do not align are extracted using SAMtools and subsequently classified into specific bacterial taxonomies using Kraken2^6^. Additionally, *BACC* includes utility scripts (utils) that enable the extraction of taxonomy-specific reads and the computation of raw count, count-per-million (CPM), and relative abundance matrices. The module generates plots of relative abundance for the most dominant bacteria within specified taxonomies across all input samples (Supplementary Fig. 1a). Additionally, phylogenetic tree with “nwk” and “tre” formats are also provided for customised visualization illustrating taxonomic relationships (Supplementary Fig. 1b).

The *BACH* module utilizes host-derived sequencing reads to predict sample-specific HLA alleles, which are expressed on the surface of tumour cells to bind and present neoantigens to T cells (Fig. 1) ^7^. Genome data is recommended for HLA typing, and *BACH* primarily targets class I HLA alleles (HLA-A, HLA-B, and HLA-C), as these are chiefly responsible for presenting intracellular antigens in tumour cells. Options for class II HLA are also provided in *BACH*. After HLA typing, the module visualizes the distribution of HLA alleles across patient samples, providing valuable insights for guiding precision medicine strategies (Supplementary Fig. 1c).

The *BACP* module predicts bacteria-derived peptides with neoantigen potentials (Fig. 1). It simulates the translational process using previously identified bacterial sequences and employs artificial neural networks and gradient-boosted decision trees trained by CheckM2^8^. Redundant amino-acid sequences are removed by CD-HIT^9^. A sliding 9-mer window is then applied to segment the effectively “translated” peptides. When proteomic data is available, *BACP* incorporates MaxQuant to analyse the proteome and annotate peptide sequences with corresponding protein IDs and associated bacterial taxonomies for further validation (Fig. 1). Subsequently, *BACP* constructs a dataset of all predicted bacterial peptides, applies the stabilized matrix method (SMM)^10^ to predict to evaluate transporter associated with antigen processing (TAP) transport efficiency, and utilizes netMHCpan 4.1^11^ to predict HLA-peptide affinity. A weight considering these two indexes is calculated as follows:

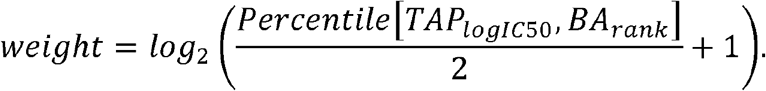

The weighted score is then utilized for peptide-HLA network visualization and prioritization, which facilitates the identification of candidate neoantigens for personalized therapeutic strategies (Supplementary Fig. 1d-1e). All three modules are integrated into a single command-line interface and can be executed independently using mutually exclusive parameters. Required dependencies are bundled within a configuration file, allowing for streamlined installation in just a few steps within an Anaconda environment.

The three modules can be performed independently or integrated together for intratumour microbiome analysis. To conduct *BACC* independently, we collected a gastric metagenome dataset (PRJNA1067082) containing 162 tissue samples. *BACC* revealed the predominance of *H. pylori*, alongside other abundant species (e.g., *S. surfactantfaciens* and *P. intermedia*) that were broadly detected in human gastrointestinal tract (Fig. 2a). Furthermore, we collected a Chinese esophagogastric junction adenocarcinoma cohort (PRJNA1067082 and PXD030667) with 98 patient samples in multi-omics data (WES, RNA-seq and mass spectrum) to comprehensively evaluate all *bacNeo* modules. After identifying bacterial taxonomies by *BACC* (Supplementary Fig. 2a), we preformed *BACH* to predict three class-I HLA alleles. In total, we identified 19 alleles in HLA-A, 38 alleles in HLA-B, and 20 alleles in HLA-C, demonstrating substantial HLA heterogeneity within the study population (Fig. 2b). Among these alleles, HLA-A11:01 was the most common one, which was consistent with previous findings in Chinese population^12^. Subsequently, we performed *BACP* which identified 3,017 unique bacterial proteins corresponding to 136 distinct species, of which 89 proteins appeared in more 20% of tumour samples (Fig. 2c). The detection frequencies of species in peptide-level and RNA-level were associated, suggesting that potential bacterial neoantigens may come from high expressed bacterial genes (Fig. 2d and Supplementary Fig. 2b). Then, *BACP* predicted bacterial peptides with high TAP transport efficiency and high HLA binding affinity, which were designated as strong binders within the analytical framework. HLA allele-specific strong binders and allele frequencies were not correlated, suggesting a diversity in binding affinities between HLA alleles and bacterial peptides (Fig. 2e). We also characterized the complicated relationships among HLA alleles, the sequences of potential bacterial neoantigens, their UniProt protein IDs, and their affiliated species, thereby presenting the most promising bacteria-derived neoantigens within this cohort (Fig. 2f). This comprehensive analysis incorporating all modules required approximately 30 minutes per sample when executed with 48 threads and up to 10 GB of memory.

**Figure 2.**
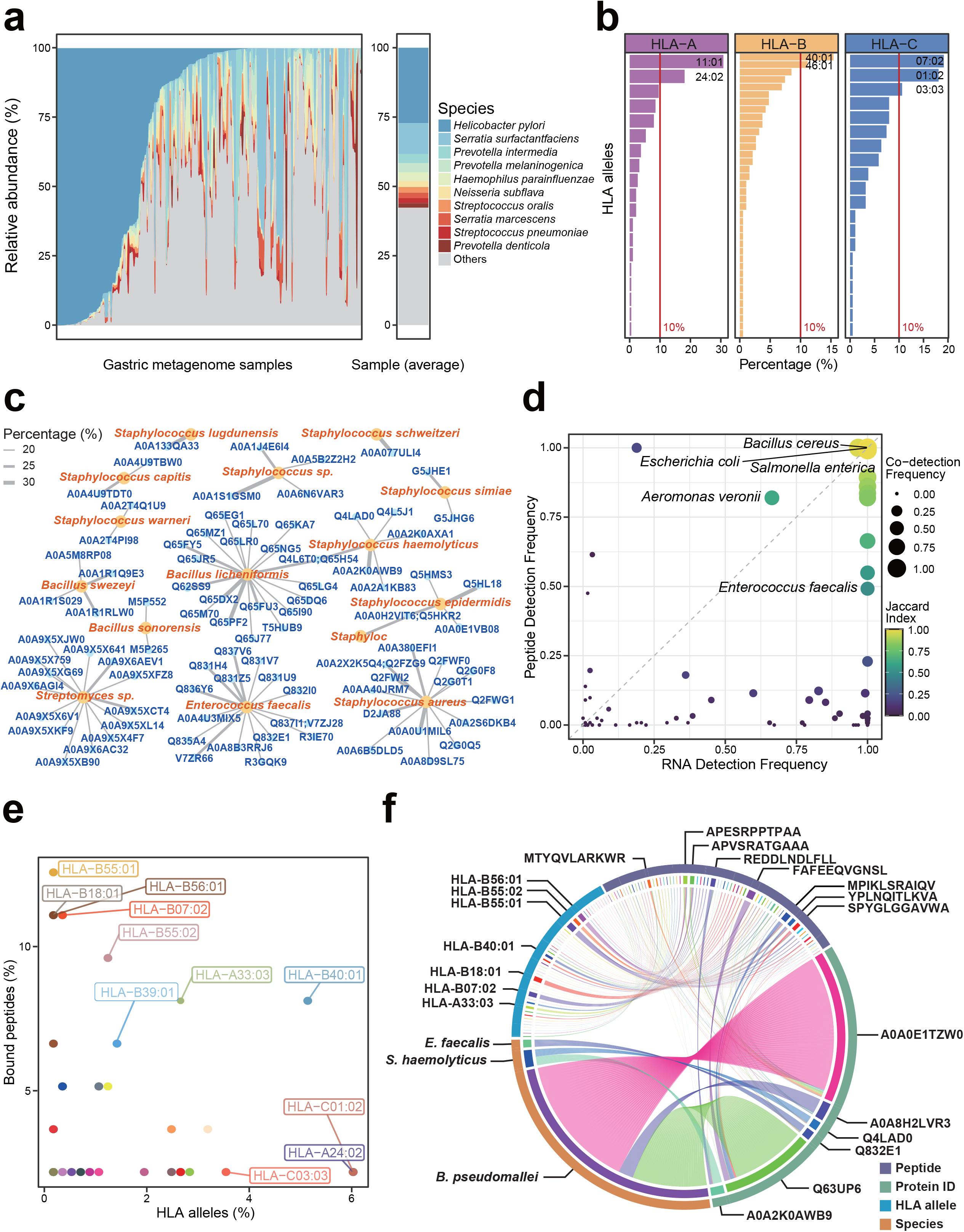
Comprehensive analysis on *bacNeo* outputs containing all three core modules. a. The distribution of abundant species identified by *BACC* in gastric metagenomic data. The samples are ordered by the relative abundance of the most pre-dominant species, *Helicobacter pylori*. Top 10 abundant species are displayed. b. The distribution of HLA alleles within class-I HLA (HLA-A, HLA-B, and HLA-C) identified by *BACP*. The red line designates 10% of total allele numbers. Alleles with frequencies above the 10% cut-off are annotated. c. The affiliation of identified peptides in species-level taxonomy. Peptides are shown in blue and annotated with UniProt Protein IDs for identification. Species taxonomy is indicated in orange with corresponding species names. Peptide abundance is displayed by the thickness of connecting edges. d. The detection frequency of bacterial species in *BACP* (y axis) and *BACC* (x axis). The point size refers to the co-detection frequency. The viridis colours refer to the Jaccard Indexes of *BACC*-*BACP* detected rates. e. The frequency of HLA alleles and their strong binders predicted by *BACP*. f. Chord plot displaying the relationships of HLA alleles, the peptide sequences of potential neoantigens, their UniProt IDs, and their affiliated bacterial species. Sections with more than ten associations are annotated with their detailed information.

In conclusion, *bacNeo* offers a streamlined, one-step *in silico* pipeline for the identification of bacteria-derived neoantigens, compatible with both single and multi-omics datasets. Its integration of machine-learning methods enables robust performance even with limited sample sizes, while multi-threading significantly improves computational efficiency. The versatility and adaptability of *bacNeo*, along with its utils, expand its potential applications and foster opportunities for open-sourced collaboration and future development. Applying *bacNeo* to single-cell and spatial transcriptomic data could further refine neoantigen discovery at higher resolution. Integration of *bacNeo* with tumour-infiltrating lymphocyte prediction tools, such as ImmuCellAI^13^ and CIBERSORT^14^, may enhance understanding of neoantigen-dependent anti-tumour effects across varying TIME. These bacterial neoantigen predictions could advance immunotherapeutic strategies—including therapeutic cancer vaccines, chimeric antigen receptor (CAR) T-cell therapy, and immune checkpoint blockade—thereby promoting precision medicine approaches across human cancers^15^.

## Online Methods

### Software practices

To test *BACC*, we initially used metagenome data from the Sequence Read Archive (SRA) database under the BioProject accession number PRJNA1067082. To test all core functional modules, we used WES and RNA-seq data from the SRA database under the BioProject accession number PRJNA788008, and proteome mass-spectrum data from in the ProteomeXchange database with dataset identifiers PXD030667^16^. The average running time for each sample is less than 30 minutes, using 48 threads and up to 10G memory. To better facilitate users for a quicker test-run, we shared a reference bash-shell script, containing three test data from BioProject (accession: PRJNA1253793).

### Reference database construction

The *bacNeo* pipeline initiates with a pre-processing module to establish reference databases. The Kraken2 database is employed for bacterial taxonomic classification. Before fully constructed, approximately 100 GB of disk space is required. A hash table would then be established for paired library sequences and taxonomic annotations, requiring approximately 29 GB of disk space^6^. The CheckM2 database containing all complete genomes in RefSeq release 202, and is integrated to assess sequence quality and filter potential contaminants^8^. These databases are automatically downloaded and formatted to ensure compatibility with downstream analyses. Pre-constructed databases can also be downloaded manually from Synapse (accession: syn66327848 and syn66514464) and put into the reference directory. In such circumstances, the construction processes can be ignored.

### Identification of bacterial reads and abundance quantification

For sequencing data alignment, BWA-MEM is utilized for genome (WGS/WES) data, and HISAT2 is applied to transcriptome (RNA-seq) data. Reads that could not aligned to the host reference genome are extracted and converted to FASTQ format using SAMtools, which are then searched against bacterial taxonomic database via kraken2^6^. These reads are mapped to different taxonomies, including domain(d), phylum(p), class(c), order(o), family(f), genus(g), and species(s). The lowercase initial(s) of taxonomic level(s) can be parsed to *bacNeo* by users based on their interested taxonomic level(s). The normalization processes would take the parsed initial(s) to form taxonomy-specific tables, including raw counts, counts per million (CPM), and relative abundance. Tables are saved under the directory of each sample.

### Construction of sample-bacteria matrices

A matrix extraction module generates sample-bacteria matrices based on user-specified taxonomic level(s) and normalization methods parsed to *bacNeo* formerly. This module further visualized abundance distributions across all samples to identify dominant bacterial taxonomies. Distribution visualisation is facilitated through the R programming language. Phylogenetic trees are also visualized in the same module, showcasing the most abundant (top 30) bacteria within user-specified taxonomic level(s). Tree visualisation is facilitated through the Python programming language, utilising functions, including NCBITaxa, TreeStyle, NodeStyle, and TextFace from package ete3^17^. A newick-formatted file (.nwk) and a tree-formatted file (.tre) are generated to the same directory, which could be input to different phylogeny platforms, for customed visualization.

### HLA allele prediction

HLAscan^7^ is implemented to predict HLA alleles from patient sequencing data. Available types include HLA-A, HLA-B, HLA-C, HLA-E, HLA-F, HLA-G, MICA, MICB, HLA-DMA, HLA-DMB, HLA-DOA, HLA-DOB, HLA-DPA1, HLA-DPB1, HLA-DQA1, HLA-DQB1, HLA-DRA, HLA-DRB1, HLA-DRB5, TAP1, and TAP2, while class I HLA types (HLA-A, HLA-B, HLA-C) are prioritized due to their role in presenting intracellular antigens, while other allele types could also be inputted to serve users’ specific needs. Predicted alleles are aggregated across samples, with a default threshold of 20% allele frequency for defining common HLA types.

### Prediction of bacterial peptides

The nucleic acid (NA) sequences of bacteria annotated in different taxonomies by kraken2^6^ are re-extracted based on utils inspired by KrakenTools^18^. Bacterial NA sequences are filtered and translated into putative amino acid (AA) sequences via CheckM2, which has been trained through artificial neural networks and gradient boosted decision trees^8^. Filtering steps are based on both the AA sequence quality and the translation potentials of bacterial peptides, which are determined by ribosomal binding motifs, start codon types, and sequence completeness. Filtered AA sequences are then segmented into 9-mer fragments, with window size of 9 and step size of 3. Redundancy reduction is performed by short word filtering algorithm integrated in CD-HIT^9^. When proteome data is provided, *bacNeo* uses MaxQuant^19^ to analyse mass spectrometry (MS) datasets to detect bacterial peptides. Detailed annotations of all peptides are generated and used as references for removing sequences homologous to humans. After all these processes have been performed, a preliminary dataset containing bacterial peptides are then put into neoantigen potential analysis.

### Prioritization of bacterial neoantigen candidates

For each peptide-HLA pair, the stabilized matrix method (SMM)^10^ is employed to predict TAP transport efficiency, as neoantigens required TAP-mediated peptide transport in endoplasmic reticulum to load onto HLA molecules^20^. This is quantified as total *TAP*_*loglC*50_ for each 9-mer peptide comprising nine amino acids, denoted as *a*(*i*) at each position *i* (where *i* ranges from 1 to 9). The *TAP*_*loglC*50_ is calculated using the following equation:

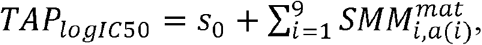

where *s*_0_ is a constant offset, and 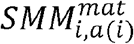 denotes the position-specific scoring matrix elements for the amino acid *a*(*i*) at position *i*^10^. The SMM matrix was previously trained to minimize the distance between predicted IC50 scores and the experimentally measured IC50 scores. Then, the netMHCpan 4.1^11^ is used to predict peptide-HLA binding affinity which is expressed as a percentile rank score, *BA*_*rank*_, computed as follows:

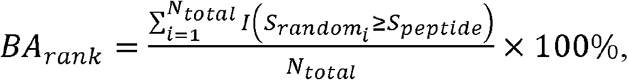

where *S*_*peptide*_ is the predicted binding score of the peptide-HLA pair, and 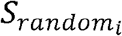 is the binding score of the *i*^th^ peptide from a large reference set of *N*_*total*_ randomly generated peptides. The lower BA_*Tank*_ indicates stronger binding affinity, and we used *BA*_*rank*_ thresholds to defined weak (below 2%) and strong (below 0.5%) binders. A weight considering these two indexes is calculated as follows:

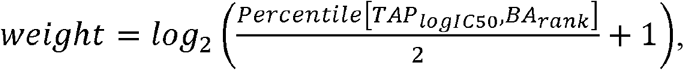

which could then be put into peptide-HLA network visualization and selection. Network visualization is facilitated through the R programming language.

### Implementation and availability

The pipeline supports multi-threaded execution, with dependencies managed via Anaconda using a YAML configuration file. Source codes and the tutorial guides are accessible on GitHub (https://github.com/ZhangLab-Fudan/bacNeo). Codes and pre-processed data for generating figures in the main figure of this article are packed within the repository for reproductivity. Users can execute modules independently or as an integrated workflow, facilitated by encapsulated scripts with detailed command-line instructions. Data for a quick test-run (SRR33242434, SRR33242436, and SRR33242438) was fetched by SRA-tools from BioProject (accession: PRJNA1253793). Pre-constructed reference databases were deposited in Synapse (accession: syn66327848 and syn66514464) to avoid potentially long waiting time in downloading and indexing processes in module 0.

## Code and data availability

*bacNeo* is open-sourced and available on GitHub (https://github.com/ZhangLab-Fudan/bacNeo), with ongoing updates and supports for issue reporting. A comprehensive user guidance and reference scripts for testing are provided in the Manual file. The test data contributing to main figures were fetched from public datasets (PRJNA1067082, PRJNA788008, and PXD030667). Additionally, a dataset containing three WES samples were fetched from public datasets (SRR33242434, SRR33242436, and SRR33242438) to facilitate quick testing for the toolkit. Pre-constructed databases are available on Synapse (syn66327848 and syn66514464).

## Acknowledgements

This research was supported by the National Natural Science Foundation of China (32270708, 82203195).

## Author contribution

Y.W. and Z.Z. conceptualized the project. Y.W. designed the methodology, analysed data, and wrote the original manuscript draft. L.Y., and X.C. provided the source data. Z.Z., K.W.G., and B.W. reviewed and edited the manuscript. Z.Z. supervised the project.

## Supplementary figure legends

**Supplementary Fig. 1. Output visualisation of *bacNeo* on the test dataset containing three samples.** a. Relative abundance distribution across samples. The plot displays the relative abundance of bacterial species at the designated taxonomic level using three test WES datasets as inputs. b. Phylogenetic tree of bacterial species generated from the Newick-format output and visualized using TreeViewer. Branches are colour-coded according to their respective phylum-level taxonomic classifications. c. Distribution of HLA alleles among patient samples, inputting the HLA-A type. Whole-exome sequencing (WES) data from all tumour samples were used as the input to determine and visualise the sample-specific HLA allele distribution. d. Scatter plot displaying both peptide-binding percentage and HLA allele percentage of each allele type. The plot illustrates the percentage of peptides predicted to bind and the corresponding frequency of each HLA allele, with HLA-A alleles selected for this test analysis. e. The predicted strong binders with top 100 weights and their binding HLA alleles. The top 100 peptide-HLA pairs, ranked by binding weights, are visualized. Weights are indicated by a colour gradient (yellow to blue) and edge thickness (thin to thick), representing increasing binding strength. Each node is annotated with the peptide’s 9-mer amino acid sequence and its associated HLA allele.

**Supplementary Fig. 2. Comprehensive analysis on the test cohort of esophagogastric junction adenocarcinoma.** a. The phylogenetic tree constructed based on *BACC*, demonstrating bacterial taxonomies in esophagogastric junction adenocarcinoma. b. The co-detection heatmap of *BACP* and *BACC*. Dark blue blocks showcase the detectable cases.

