## Supplementary figures and images for "bacNeo: a computational toolkit for bacteria-derived neoantigen identification"

### Supplementary Figure 1

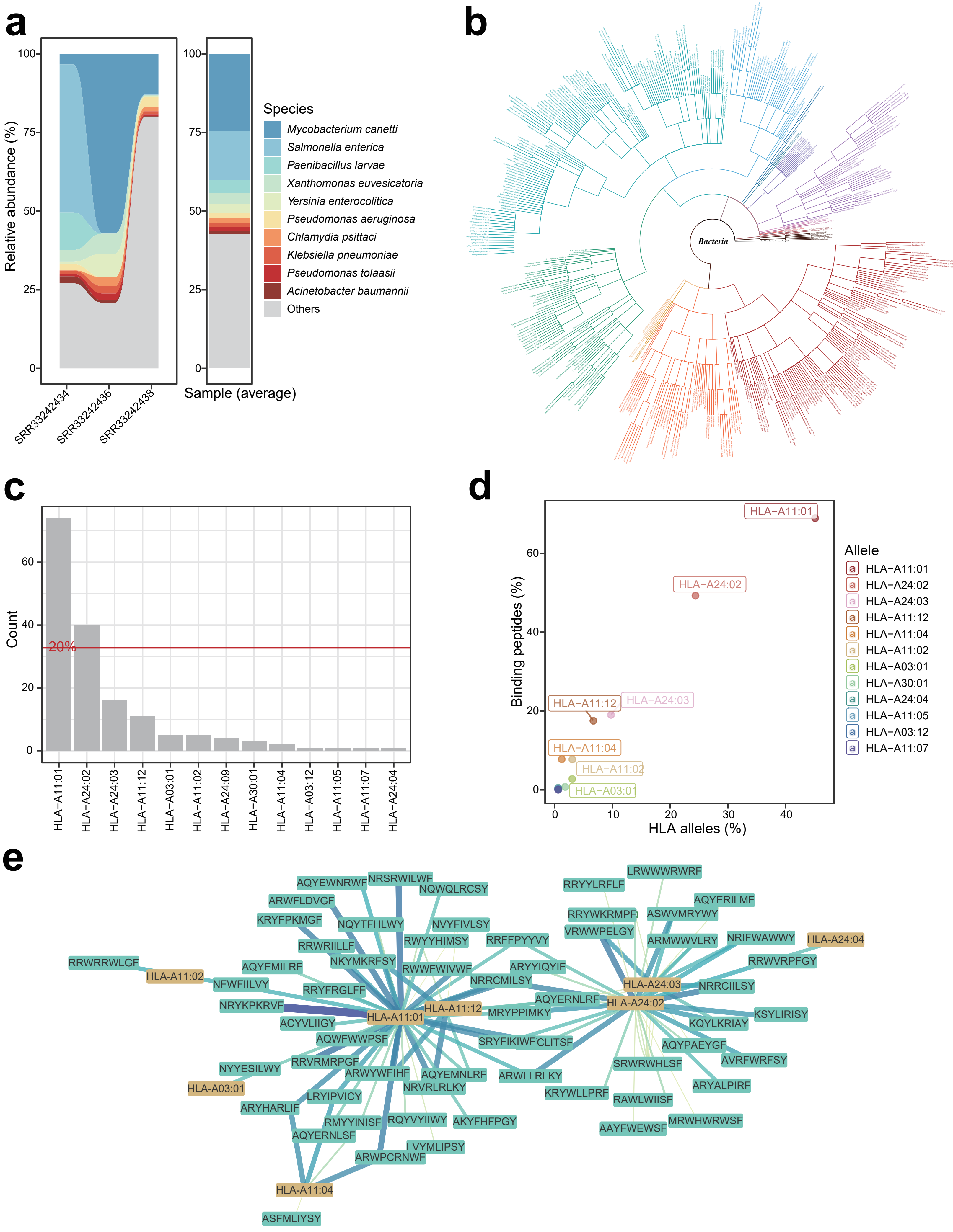

### Supplementary Figure 2

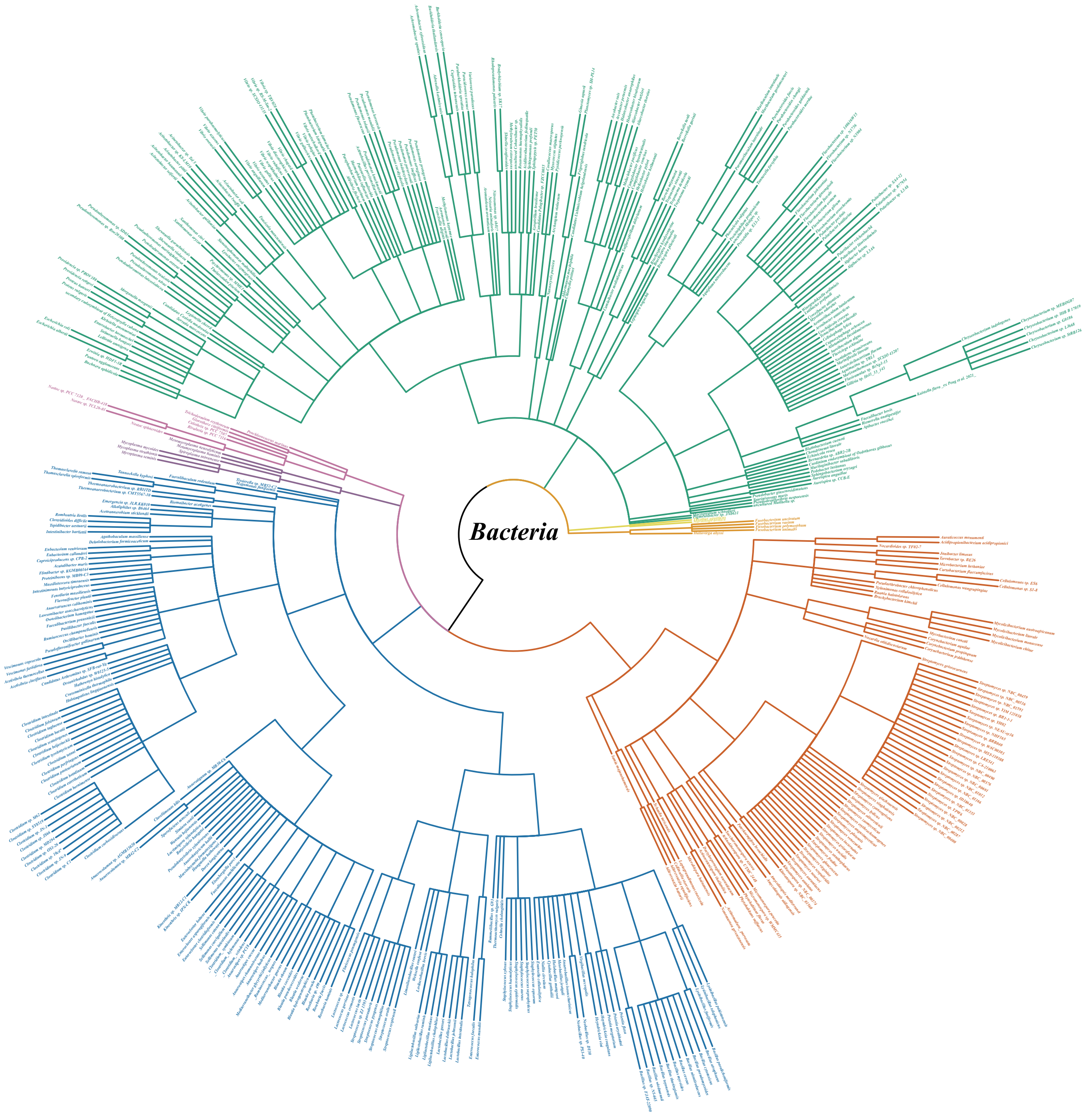

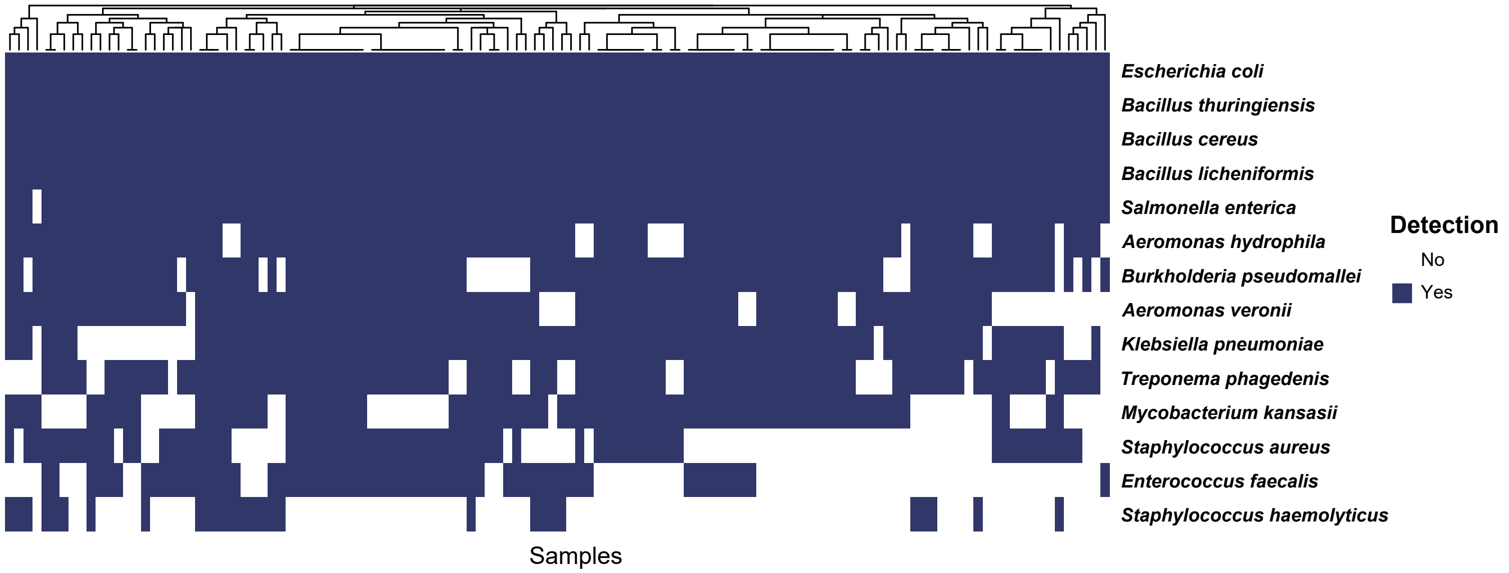
